# *TiFoSi*: an Efficient Tool for Mechanobiology Simulations of Epithelia

**DOI:** 10.1101/2020.02.12.933903

**Authors:** Oriol Canela-Xandri, Samira Anbari, Javier Buceta

## Abstract

**About:** This document is an extended version of the main text where some details and results are fleshed out. Further details can be also found in the manual of the code and at *TiFoSi*’s website: http://tifosi.thesimbiosys.com.

**Motivation:** Emerging phenomena in developmental biology and tissue engineering are the result of feedbacks between gene expression and cell biomechanics. In that context, *in silico* experiments are a powerful tool to understand fundamental mechanisms and to formulate and test hypotheses.

**Results:** Here we present *TiFoSi*, a computational tool to simulate the cellular dynamics of planar epithelia. *TiFoSi* allows to model feedbacks between cellular mechanics and gene expression (either in a deterministic or a stochastic way), the interaction between different cell populations, the custom design of the cell cycle and cleavage properties, the protein number partitioning upon cell division, and the modeling of cell communication (juxtacrine and paracrine signalling). *TiFoSi* fills a niche in the field of software solutions to simulate the mechanobiology of epithelia because of its functionalities, computational efficiency, and its user-friendly approach to design *in silico* experiments using XML configuration files.

**Availability:** http://tifosi.thesimbiosys.com

## 1 Introduction

The interplay between mechanical and biochemical cues drives cellular behavior and determines features such as proliferation, motility, and differentiation. Cell interactions further scale up the mechano-signalling feedbacks and regulate key elements of the collective behavior such as patterning and the control of size and shape in tissues. All these questions lie within the interest of the field of mechanobiology (Jacobs *et al*., 2012). Importantly, a number of pathologies, such as cancer progression (Carey *et al*., 2012), depend on a misregulation of mechano-signalling cues. Moreover, fundamental open questions in the field of developmental biology are rooted in the feedback between gene regulation and mechanical responses. For example, keeping cellular populations segregated is essential for the growth, patterning, and the formation of tissues/organs during development and it has been shown that this phenomenon relies on the interplay between physical mechanisms (e.g., increased cell bond tension) and gene expression (Aliee *et al*., 2012). In this context, there is a growing role for computer simulations of tissues because they assist in the quantitative understanding of the experimental phenomenon and help to test/suggest hypotheses (Schwarz and Dunlop, 2012; Morelli *et al*., 2012).

Surprisingly, while the amount of simulation tools available to researchers for other biology-related problems is overwhelming (e.g., simulation of gene regulatory networks or bioinformatics suites), in the case of tissues the offer of software packages is very limited. This is mainly because 1) tissue simulations are intrinsically multi-scale covering time scales from seconds (e.g., cellular mechanics) to days (e.g., tissue remodeling) and, consequently, complex, 2) they are context dependent and thus require tailored solutions, and 3) the possibility of including feedbacks between the cellular mechanical properties and biochemical signals (e.g., gene regulation) represents an additional challenge in terms of the computational efficiency. As of today, to the best of our knowledge, there are just two multi-scale tissue simulation tools publicly available to researchers that are able to deal, up to some degree, with mechanobiology problems. On the one hand, CompuCell3D (CC3D) is based on the cellular Potts model (Graner and Glazier, 1992). This software poses several advantages: it is well-documented, has a good user support through an online forum, and it is scriptable (Swat *et al*., 2012). However, CC3D has also some disadvantages: the simulation methodology is based on a Montecarlo approach that makes it difficult to connect time steps with real time in experiments. More importantly, CC3D is based on an on-lattice method where each cell is defined by a collection of lattice points. Thus, the simulation of large tissues is very costly, computationally speaking. This problem has been acknowledged, and code parallelization solutions were proposed to scale-up the capabilities of the software (Chen *et al*., 2007); yet, that solution is not fully implemented and, in any case, only access to computer clusters can improve CC3D efficiency when dealing with large tissues. On the other hand, Chaste (Cancer, Heart and Soft Tissue Environment) is a C++ generic extensible library aimed at multi-scale, computationally demanding problems arising in biology (Mirams *et al*., 2013; Pitt-Francis *et al*., 2009). In particular, Chaste allows to implement simulation of tissues using the vertex model methodology (Fletcher *et al*., 2013, 2014). The vertex approach is off-lattice and, crucially, it is computationally efficient since cells are represented by a small set of points (the vertexes that define their shapes). For example, a simulation of a tissue with 10^3^ cells would require to simulate the dynamics of 𝒪 (10^5^) points using the Potts model but only 𝒪 (10^3^) points using a vertex model approach (i.e., a two orders of magnitude improvement). Our own experience implementing simulations using the vertex model reveals that regular PCs can deal efficiently with tissues with a large number of cells (Canela-Xandri *et al*., 2011; Buceta, 2017). On the downside, Chaste is not a user-friendly software, but a collection of libraries that require C++ programming knowledge in order to setup and run a simulation. Thus, this restricts its usability. Altogether, there is a gap of development that needs to be filled in regards to multi-scale tissue simulation tools.

Here we fill that gap by introducing *TiFoSi* (*Ti*ssues *Fo*rces & *Si*gnaling). This simulation software includes the advantages of the tools mentioned above and avoids their weaknesses. Specifically, it uses a vertex model methodology, it is user-friendly by using a scriptable approach (XML), yet computationally efficient (C++). In addition, *TiFoSi* is open, fully documented, and data-centric (see below). We expect that *TiFoSi* will increase the usage of computational approaches in research groups in the field of tissue dynamics, including mechanobiology, with a limited computational background.

## 2 Methods

### 2.1 Design and Implementation

The design and implementation of *TiFoSi* adhere to the following guidelines:

- **Fast and efficient**. The complexity of tissue simulations makes interpreted languages highly inefficient in terms of execution speed. Thus, the code must be based on a high-end compiled language (in the case of *TiFoSi* C++).
- **User friendly**. Previous efficient software solutions for simulating epithelial dynamics following a vertex model approach rely on the knowledge of a high-end computer language (i.e., C++). In order to circumvent this major disadvantage, *TiFoSi* relies on XML configuration files with a syntax and modularity aimed at clearly defining *in silico* experiments. A Python parser then interprets the configuration file and modifies the C++ source code. The learning curve to create configuration files is fast and we provide an extensive manual that explains all the possibilities of the code with examples.
- **Open**. *TiFoSi* uses free open source languages (GNU g++ and Python 2.7) and the code is available under a GNU license. *TiFoSi* has been extensively tested in Linux OS (Ubuntu) but will run in any platform where GNU g++ and Python 2.7 are installed.
- **Data oriented**. Tissue simulations present endless possibilities in terms of the conceivable experiments. Thus, focusing on the development of a graphical frontend, both for the input (configuration file) and the output, would make the code extremely complex, would pose problems in terms of its platform portability, and would make the code less flexible for implementing some functionalities. For these reasons, *TiFoSi* input and output are data oriented. On the one hand, as mentioned above, input is based on an XML configuration file. On the other hand, *TiFoSi* produces a comprehensive collection of output files and logs (ANSI text) that report on all the possible relevant information of the tissue simulation including cells geometrical properties, energy, acting forces, protein expression levels, cell lineage tracking information, and properties of division events and tissue topology changes.
- **Community Sustained**. We aim at developing a community of *TiFoSi* users and developers. *TiFoSi*’s website (http://tifosi.thesimbiosys.com) is linked to the OSF (Open Science Framework) project’s website to ensure long-term support and users’ collaborative efforts. Thus, the project’s site has been conceived as a repository of configuration files, visualization tools, further modifications of the code, and, importantly, a forum to discuss issues and questions among users and developers.

### 2.2 The underlying modeling approach: modularity

On top of sections focused on the definition of the initial conditions and the possible different stages of a simulation, the configuration files in *TiFoSi* distinguish three main modules to define an *in silico* experiment: cell mechanics, cell growth/cleavage, and protein dynamics.

#### Energetics: the vertex model

In the cell mechanics module, the parameters that define the mechanical properties of the cell are specified. Thus, following a vertex-model approach, every cell in the tissue is represented by a discrete set of points: the vertexes that define the polygonal shape of the apical surface of epithelial cells. In *TiFoSi* the energy functional acting on a cell vertex *i* reads,

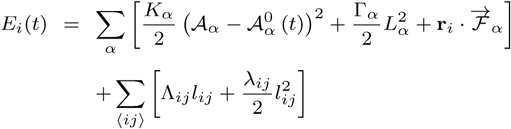

where the sums indexed by *α* and ⟨*ij*⟩ run respectively over the cells, *α*, and the vertices, *j*, sharing vertex *i*. The terms on the right hand side represent different energetic contributions. The first term accounts for the elastic energy of cells, *K*_*α*_ being proportional to the Young modulus, due to the difference between the *actual* cell area 𝒜_*α*_ and the one that the cell would have due to the cytoskeleton structure (but in the absence of the stresses associated to adhesion and cortical tension), 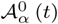 (*t*). The Λ_*ij*_ term stands for the line tension, *l*_*ij*_ being the length of the edge connecting neighboring vertices *i* and *j*, and includes contributions from cell-cell affinities (i.e., the action of molecules regulating the adhesion between cells) and also the cortical tension. The Γ_*α*_ term mimics further contributions from the tension associated with the contractility of the actomyosin ring. This term is proportional to the squared cell perimeter, *L*_*α*_, and takes into account a global contractility effect. The *λ*_*ij*_ term accounts for possible inhomogeneities of the contractile tension due to the accumulation of actin-myosin in specific regions of the ring. Finally, the 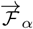 term (**r**_*i*_ being the position of vertex *i*) accounts for the effect of additional forces acting of the cell mimicking either external (e.g., pulling) or internal (e.g., active migration) forces. In *TiFoSi* all the mechanical parameters (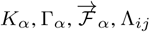), and λ_*ij*_ can be dependent on gene expression levels, the cell cycle properties, the cell type, etc.

By realistically neglecting inertial effects with respect to dissipation, tissues are driven by an over-damped dynamics with a characteristic kinetic coefficient *γ*. Thus, the equation of motion for vertex *i* at position **r**_*i*_ driven by the conservative force **F**_*i*_ = −∇*E*_*i*_ may be written as,

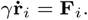

In *TiFoSi* this equation is numerically integrated using an Euler algorithm, and vertexes evolve off-lattice. Further details about the dimensionless form of the energy functional and the vertex recombining processes that provide tissue plasticity, e.g., T1 transitions, can be found in the manual (Supplementary Information).

#### Cell cycle dynamics: tissue growth

The cell cycle module in the configuration file defines the growth and division properties of cells. In that regard, the cell cycle duration is not deterministic but stochastic and depends on cell-autonomous processes and the mechanical interactions with the local cellular environment. Both effects are considered in *TiFoSi*. Moreover, during the course of the cell cycle, distinct growth phases can be identified. We account for these observations as follows. As for the cell cycle duration, we have defined an internal “clock” for each cell that measures the time elapsed since the beginning of the cell cycle after division. The variable *τ*,

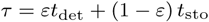

accounts for the cell cycle duration in a cell such that *t*_det_ is a deterministic time scale that accounts for a mean cell cycle duration (in the absence of mechanical stress due to the local cell environment), and *t*_sto_ is a random variable that accounts for its variability. This last contribution, *t*_sto_, is assumed to follow an exponential distribution:

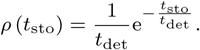

The parameter *ε* ∈ [0, 1] controls the relative weight of the deterministic and stochastic components such that, with the above definitions, the average and standard deviations of *τ* are given respectively by,

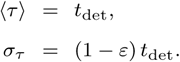

As for the different growth phases, those are implemented in the code by prescribing a particular dynamics for the *preferred* apical cell area 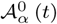. The *actual* duration of the cell cycle, that leads to the division event, depends on conditions set over such dynamics as detailed in the manual (e.g., cell side doubling).

With respect to the cleavage orientation, the code evaluates the inertia tensor of the cell with respect to its center of mass (assuming that a proper representation of the cell is a polygonal set of rods, i.e., the cell edges). Upon diagonalization of the inertia tensor the principal inertia axes are computed and subsequently the longest axis of the cell (orthogonal to the direction along the largest principal inertia axis). This allows to define a criterion for the cleavage orientation depending on the cell geometry, e.g., the cells may divide satisfying the Hertwig rule (cleavage plane perpendicular to the longest cell axis), or opposite to the Hertwig rule, or randomly, since *TiFoSi* also allows to include variability in the cell cleavage orientation. Following cell cleavage, proteins (or any other species, e.g., plasmids) are distributed among daughter cells, either equally (i.e., half-and-half) or satisfying a binomial distribution depending on the user input in the configuration file.

#### Gene Regulation: Protein Dynamics

This module simulates the dynamics of molecules within the cell. We stress that although we use the term “protein” (hereafter and in the manual), any species or quantity of interest that evolves dynamically can be simulated (e.g., ribosome content, cell identities, number of neighbors…). In this regard, *TiFoSi* assumes that spatial effects within the cells can be disregarded (i.e., cells are considered well-stirred compartments).

The dynamics is then prescribed by specifying a growth rate for the *number* content of proteins. Thus, given a set of proteins {*X*_*j*_} = {*X*_1_, *X*_2_, …, *X*_*N*_} such that their dynamics is given by the differential equations 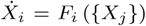, the growth rate functions *F*_*i*_ ({*X*_*j*_}) must be specified in the code. *TiFoSi* includes a small library of functions commonly used to model gene regulation, such as the Hill functions, white noise terms for stochastic simulations, and functions to prescribe juxtacrine and paracrine signalling. Further details about the library of predefined functions are provided in the manual. The algorithm used in the code for the numerical integration of the equations is Euler. While smaller errors could be achieved with other simple, explicit, integration schemes (e.g., Heun algorithm), the main advantage of the Euler scheme is the integration of stochastic differential equations since it converges to the so-called Ito calculus interpretation (that is the one with a physical meaning in the context of stochastic simulations of protein expression due to low copy number (Canela-Xandri *et al*., 2010)). Also, we notice that the maximum accuracy that can be obtained when integrating stochastic differential equations using explicit methods is Δ*t*^3/2^, Δ*t* being the time step, regardless of the order of the integration scheme (Euler accuracy being of order Δ*t* (Rümelin, 1982)).

In summary, three different modules define a tissue simulation implemented in *TiFoSi*: the tissue mechanics, the cell growth/cleavage, and the species dynamics (e.g., gene regulatory processes). As detailed in the manual, and illustrated by the example below (see Results), the code allows to include dependencies between them to implement *in silico* mechanobiology experiments. E.g., the cell growth or the adhesion between cells can be dependent on the expression levels of proteins.

### 2.3 Output/Input files and logs

As mentioned above, *TiFoSi* is data-centric and provides a comprehensive output in terms of regular text data files:

- **dcells**.**dat**: data about cell identities for tracking purposes, cellular geometric properties (area, cell center, and coordinates of the vertexes), information about neighboring cells, and “protein” expression levels (number).
- **dforces**.**dat**: data about the value of the different forces acting on each cell vertex.
- **denergy**.**dat**: data about the value of different energetic contributions in each cell vertex.
- **divisions**.**dat**: data about cell division events including mother and daughter cells identities, timing, orientation and other geometric information relative to the cell cleavage process.

Additional log files provide data about tissue topological changes (e.g., T1, T2, and T3 transitions, see manual), changes (if any) of the value of the parameters of the energy functional as time progresses, list of the proteins involved in the simulation, last frame of every temporal stage during the simulation, and a simulation progress log file that indicates the time consumed for simulating each frame. Details about the structure of data files and logs can be found in the manual.

Finally, *TiFoSi* also allows to import data from previous simulations or from experiments assuming that a segmentation has been performed and the data is provided following the format of the output file dcells.dat.

## 3 Results

In order to illustrate some of the capabilities and the syntax of *TiFoSi*’s configuration files, we provide simulation results of a case study: formation and remodeling of Turing patterns in growing epithelia. In that regard, during the last decades different studies have shown the widespread biological applicability of this patterning mechanism in development (Kondo and Asai, 1995; Kondo and Miura, 2010; Raspopovic *et al*., 2014; Hentschel *et al*., 2004; Miura *et al*., 2006; Murray, 2001; Jernvall *et al*., 2010; Economou *et al*., 2012). We stress that the example case presented here does not necessarily apply to a particular biological problem and just aims at showing the main features of the simulation tool and how to implement mechanobiology feedbacks.

Thus, we implement a generic protein-protein reaction-diffusion scheme that undergoes a Turing’s instability (Chen and Buceta, 2019):

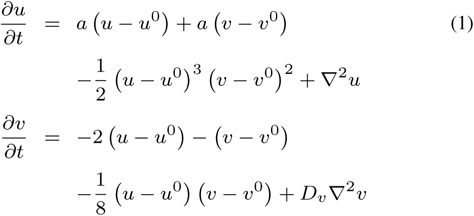

In eqns. (1), *u* and *v* represents the dimensionless cellular *concentrations* of an activator and an inhibitor respectively. Under the following conditions a pattern develops (Chen and Buceta, 2019),

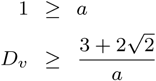

In our simulation we use the following parameters: *a* = 0.9, *D*_*v*_ = 9, *u*_0_ = *v*_0_ = 2.

The simulation is divided in two stages that reveal how tissue growth induces pattern remodeling and how mechanobiology feedbacks can modify growth. During the first stage, tissue growth and protein expression are implemented but decoupled. During the second stage, the cortex activity (contractility) depends on the expression levels of the species *u* and *v* and induces additional pattern remodeling and shape changes at the tissue level (see Fig. 1).

**Fig. 1.**
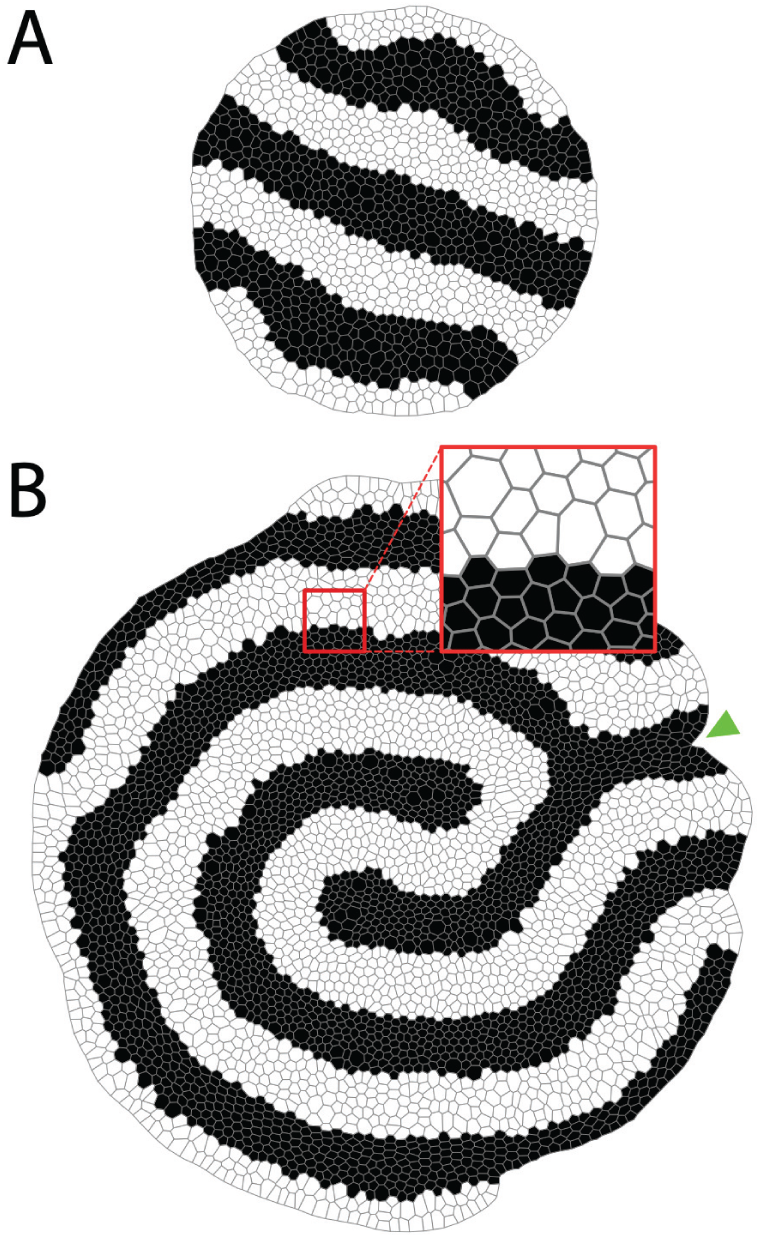
**A** and **B** panels show snapshots at the end of the two temporal stages defined to implement the simulation of the Turing patterning signalling. Black: *u* > *v*; White: *v* > *u*. During the first stage, signalling and mechanics are decoupled and patterning is simply conditioned by tissue growth. During the second stage, cells regulate their cortex activity dynamically depending on their patterning domain: as revealed by their size (inset), cells in domains where *u* > *v* have larger contractility than cells in domains where *v* > *u*. These mechanical differences can induce tissue remodeling effects such as indentations (green arrow)

In the examples’ section of *TiFoSi*’s website (*tifosi*.*thesimbiosys*.*com*) we provide the XML configuration file for performing this simulation together with all the generated output files and also movies obtained by processing these files using a Wolfram’s Mathematica notebook (available at the Visualization Tools section). As revealed by the log file time.dat, the total execution time for performing this simulation showing a cellular growth from 10^1^ to ∼ 4 · 10^4^ cells (an average of ∼ 5 cell cycles per cell) was ∼ 6 minutes using a computer equipped with an Intel Xeon CPU E3-1220 v5 3.0GHz and 16Gb RAM (Ubuntu Linux v18.04.3, g++ v7.4.0, and Python v2.7.15+).

Figure 1 shows tissue snapshots at the end of each simulation stage. Further details about the patterning and tissue remodeling (including cell division events) are noticeable in the simulation movies.

## 4 Discussion

Here we have introduced *TiFoSi*, a tool to perform versatile tissue simulations. *TiFoSi* is based on the vertex model methodology and is computationally efficient and user-friendly by using a scriptable approach to design *in silico* experiments. Importantly, *TiFoSi* allows to couple cell mechanics, gene regulation, and cell cycle/cleavage properties, thus enabling mechanobiology simulations. As for the limitations of this software, this tool is useful in the context of planar epithelia and focuses on the dynamics of the apical side of cells. Yet, recent results have shown that the cellular dynamics in epithelia when subjected to bending and folding requires further considerations since apico-basal intercalations (*scutoids*) develop (Gómez-Gálvez *et al*., 2018). Thus, possible extensions and future directions include a generalization to a 3D context to realistically consider the bending and folding effects in the cellular geometry.

As a matter of discussion, the promising field of organoids offers an exceptional framework to study developmental biology and personalize medicine (Ader and Tanaka, 2014). While 3D culturing is a key component for the creation of organoids, some problems, such as the causes and consequences of their natural variation (lack of reproducibility), can be explored by simplified models in 2D. Also, there is growing evidence that mechanobiology pathways and physical forces are key elements to understand organoids dynamics, e.g., (Martinez-Morales *et al*., 2017). Given that *TiFoSi* allows stochastic modeling of tissues that comprise multiple cell populations with distinct mechanical properties, we argue that this tool can be useful to move the field forward.

Altogether, we believe that *TiFoSi* fills a niche in the field of software solutions to simulate the mechanobiology of tissues and that because of its functionalities, efficiency, and user-friendly approach will promote the usage of computational approaches in research groups with a limited computational background and will help to move the field of developmental biology, including organoids research, forward.

## Funding

This work has been supported by Lehigh University through a Faculty Innovation Grant (FIG-2019-JB).

## Notes

http://tifosi.thesimbiosys.com

